# Confirmation of fishers’ local ecological knowledge of ciguatoxic fish species and ciguatera-prone hotspot areas in Puerto Rico using harmful benthic algae surveys and fish toxicity testing

**DOI:** 10.1101/2021.10.10.463731

**Authors:** Henry Raab, Joe Luczkovich, Miguel Del Pozo, David Griffith, Cindy Grace-McCaskey, Wayne Litaker

## Abstract

Ciguatoxin fish poisoning (CFP) is caused by the consumption of tropical and subtropical fishes and other marine species with high levels of ciguatoxin (CTX) in their tissues. CTX is a polycyclic neurotoxin produced by single-celled, photosynthetic dinoflagellates in the *Gambierdiscus* and *Fukuyoa* genera which are found in close association with benthic autotrophs. CTX enters the food web when these dinoflagellates are inadvertently consumed by herbivores grazing on their preferred substrates. The toxin biomagnifies up the food chain to the top predators and if humans consume seafood with high levels of CTX it can cause a variety of flu-like symptoms. The best way to avoid CFP is to avoid toxic fishes. However, CTX is undetectable by physical inspection. This study investigated local fishers’ knowledge of ciguatera hotspots and coldspots along Puerto Rican coral reefs using toxic-dinoflagellate cell counts and by estimating fish toxicity in those sites using a cell-based Neuro-2a cytotoxicity assay. The fishers identified regions of high and low risk for CFP based on their local ecological knowledge (LEK) which were deemed hotspots and coldspots, respectively. There is a 35-fold difference in dinoflagellate cell counts of low-toxicity G*ambierdiscus* species in samples in the identified hotspot compared to the coldspot. Also, higher trophic level fishes (>3.4 ETL) had higher median estimates of CTX in their tissues at the hotspot than the same species in the coldspot. This study shows the effectiveness of LEK in identifying potential problem areas for ciguatera.

## Introduction

People living in tropical and subtropical regions worldwide rely on fish and other marine organisms for sustenance, tourism, and recreation. However, fishes in these regions, specifically in the Pacific and Indian Oceans and the Caribbean Sea, can harbor ciguatera toxin (ciguatoxin or CTX), a potent neurotoxin produced by several different species of dinoflagellates, most notably in the *Gambierdiscus* and *Fukuyoa* genera [1,2]. If humans ingest tissues of marine coral reef species that accumulate this toxin in a high concentration, then it can cause a variety of severe symptoms, i.e., vomiting, diarrhea, abdominal pain, paresthesia (burning of the skin), the reversal of hot and cold sensations, and occasionally, death [3]. The muscle tissues (the fish filets most people consume) have the potential to be toxic. Also, the roe, gonads, liver, and other organs in the fishes carry higher levels of CTX than muscle tissues, and these organs may be more dangerous to consume than muscles [4]. Different structures and chemical congeners of ciguatoxins in the Indian Ocean, Pacific Ocean, and the Caribbean Sea cause variations in symptoms from those regions [5–7]. The sickness from consuming ciguatoxic fish is known as ciguatoxin fish poisoning (CFP).

There is no reliable way to determine if seafood has high levels of CTX, the best way to prevent CFP is to avoid it altogether. CTX is colorless, odorless, and tasteless [8] and is heat-stable, meaning cooking the fish does not affect the toxin [7]. Local folk methods for identifying toxic fish (such as feeding a small piece of fish to a pet animal and monitoring its reaction, rubbing the flesh with a coin, or leaving a portion of the fish near insects to see if they avoid it) are unreliable [9]. There are currently no dockside tests available. The best ways to detect CTX are with complex bioassays (neuroblastoma cell-based assay (N2a-CBA), and a fluorescent receptor binding assay (RBA(F))) which are costly and time-consuming [10–12].

Local ecological knowledge (LEK), also known as traditional ecological knowledge (TEK) [13] is the knowledge and beliefs about ecological relationships gained from interaction with a resource that can be shared among other resource users [14]. It has been shown that indigenous fishers can understand fishes' migration patterns, habitat connectivity, population dynamics, essential fish habitat, and the presence or absence of species [15–17]. Therefore, we believe that Puerto Rican fishers can identify reefs that have high and low levels of ciguatoxin, which we are calling “hotspots” and “cold spots”, respectively. Our theory is that over time, fishers have learned to avoid certain reefs, and fishes within those reefs, due to a feedback loop where the harvesting and consumption of toxic species changed fishing habits to avoid these toxic areas and species. We show here that reefs identified by fishers as hotspots had 35-fold more toxin-producing dinoflagellates than areas identified as cold spots. There was also more CTX in the fishes’ tissues in the higher trophic levels (>3.4 ETL) at the hotspots than those same species in the coldspots using CTX estimation with the reliable N2a-CBA neuroblastoma cell-based assay.

## Materials and Methods

### Interviews with Fishers

Semi-structured interviews were conducted with 21 commercial fishers in Puerto Rico to identify hotspot and coldspot locations to sample fishes for CTX estimation and determine which fishes would likely have higher levels of CTX in those areas. These data guided the protocol for both the fish and toxic dinoflagellate sampling. Interviews took place in *Villas pesqueras* (fish houses) along the west, south, and east coasts of Puerto Rico. The informants’ commercial fishing experience ranged from 18 to 67 years 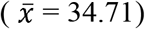, thus they possessed a great deal of ecological knowledge on fishes harvested commercially, coral reefs, and ciguatoxic fishes. Informants circled areas they identified as hotspots and coldspots on nautical charts (NOAA booklet charts 25650, 25977, and 25668). The closest municipality to the circled area's location was designated as the name of that hotspot or coldspot. For example, a circled area off the coast of Guayama was simply “Guayama.” Each fisher had a new booklet chart to draw on to discourage biased results from previous fishers. Hotspots and coldspots were identified using a free-list salience metric in Visual Anthropac [18]. Salience is a weighted average of the inverse rank of an item across multiple free lists, where each list is weighted by the number of items in the list [19]. The salience metric was used to analyze the names of the high-prevalence CFP locations in the free lists provided by the respondents and to compare the different fishing villages in terms of their ranking of locations for CFP. Salience was also used to compare three larger coast-wide regions (Northeast coast, South coast, and Southwest coast).

The informants also identified fishes they believed to be at higher risk of CTX. As part of the interviews, we asked the fishers to free-list species that they consider to be toxic. These free-lists were analyzed using Visual Anthropac using the salience metric as described above for CFP locations. After free-listing we asked them to sort fish cards into two piles: toxic and non-toxic. This pile sort exercise was done with a fixed set of fish species identification cards that was administered to all interviewees to investigate which fishes and other marine species they believed were most toxic. These data were used to identify which fishes would be best for sampling. A set of laminated cards were created with different species of fish on each one. The fishes on the cards consisted of commonly caught species of commercial value in Puerto Rico except for *Sphyraena barracuda*. Barracuda were included because they are known to have high levels of CTX in their tissues, and the Puerto Rican government has a moratorium on the commercial catch and sale of this species due to CTX concerns. The informants put the cards into two piles according to whether they avoid catching that species due to CTX or not. Results were analyzed using a consensus analysis in UCINET [20]. Consensus analysis examines a respondent matrix, with informants as rows *i* and fish species cards as columns *j,* with cells coded with a 1 if placed by an informant in the toxic pile, and 0 if in the non-toxic pile. This two-mode adjacency matrix was analyzed with the UCINET Consensus Analysis tool and plotted in NetDraw [21]. Consensus analysis determines the most likely “correct answer” amongst the respondents and simultaneously assesses the competence of each of the respondents. We formatted the pile sort data as a matrix with fishers as respondents and the response for each species as an answer to a question: “Is the species toxic? True or False”. Each fish species card was placed by the respondent in a True or False pile. The fisher true-false responses for each species were recorded in a matrix, with 1=True and 0=False. The consensus analysis produced an agreement matrix of the fishers’ responses, a square data matrix that indicated the consensus among fishers. Also, the consensus analysis produced a “correct” answer key for each species of fish, indicating which species most fishers agreed were toxic. The consensus analysis routine produced a competency score for each fisher, and output of eigenvalues, with the Eigenratio (ratio of first to the second eigenvalue) providing a reliability estimate for the analysis.

### Toxic Dinoflagellate Sampling

Based on LEK interviews toxic dinoflagellates were sampled in October 2019 at four sites: CTX-1 (23.7m to 25m depth) and CTX-2 (18.9m to 21.6m depth) were identified as coldspots and CTX-3 (22.5m to 28.6m depth) and CTX-4 (17.2m to 18.9m depth) were identified as hotspots. We used the artificial substrate screen-rig 24-h sampling protocol outlined in Tester et al 2014 [22] to provide consistent substrate area and type across these four sites. Five replicate screen-rigs were deployed on the open bottom away from coral heads on open or algal-covered substrates at each site by scuba divers, with one central screen-rig and four others surrounding this central one spaced at distances of 10-45m apart. The rigs were a simple weight attached to a fishing bobber with a barrel swivel attached 1m from the weight and a mesh screen attached to a swivel. After 24 hours, divers collected the rigs by placing a jar over the screens and unhooking the swivels; the lids were tightened on the jars and brought to the surface. The samples were taken to the University of Puerto Rico at Humacao and preserved with Lugol’s solution. The water samples were stored in brown plastic bottles and then shipped to the NOAA Southeast Fisheries Science Center. The samples were counted for the number of *Gambierdiscus* spp. cells present using a Nikon Eclipse TS100 at 40x magnification. Species were identified in each sample using a semi-quantitative polymerase chain reaction (qPCR) [23,24].

### CTX Estimation of Fishes at Hotspots and Coldspots

We sampled fishes at two reefs at two different depths in October 2019 for two consecutive days at the identified coldspot and the hotspot. Fishes of all trophic levels were targets for the study; however, hogfish (*Lachnolaimus maximus)* and barracuda (*Sphyraena barracuda)* were a high priority. Barracuda were targeted due to their high trophic position (~4.0) in the food web and the commercial harvesting ban of these fishes. Hogfish were targeted because of their trophic position (~3.66) and they are a commercially important species to the fishers of Puerto Rico that have been known to harbor toxic levels of CTX. Some informants mentioned them as highly toxic in some areas and non-toxic in others. The tested fishes were captured by local fishers by scuba diving, fish traps, and hook and line. For analysis, fishes were separated into three trophic groups, based on their ETL’s from Opitz (1996) [25]. The low trophic group was 2.0 – 2.9 ETL, the medium trophic group was 3.0-3.4, and the high trophic group was 3.5+. This was based on diet compositions. The low trophic group was mostly herbivorous with some zooplankton. The medium trophic group consumed a mixture of plants, invertebrates, and fishes, and the high trophic group ate mostly fishes. The low trophic group solely consists of *Sparisoma viride*, the medium trophic group was *Holocentrus rufus,* and *L. maximus,* and the high trophic group was *S. barracuda* and *C. ruber*.

Muscle tissue was taken and stored from each fish caught which were used in the N2A-CBA neuroblastoma cell-based assay to estimate CTX concentrations. First, CTX was isolated from the muscle tissues and suspended in 100% methanol. Five grams of fish tissue were homogenized twice in 10ml 100% methanol in a 50ml Falcon centrifuge tube using an electric tissue homogenizer. After each homogenization step, the methanol was transferred from the 50ml Falcon tube to a glass HPLC scintillation vial. It was essential to use glass vials because CTX can stick to plastics. The methanol layer was allowed to dry under an N^2^ stream until only the precipitate remained. Then, 5ml dichloromethane (DCM) and 5ml 60% methanol were added to the glass scintillation vial twice.

After each substance’s addition, the vial was swirled, then its contents were added to a 250ml glass separatory funnel. The layers were separated after shaking lightly, and the DCM layer was added in a new glass scintillation vial. The N^2^ stream dried the sample until the precipitate remained. Next, 5ml cyclohexane and 5ml 80% methanol were added to the new glass scintillation vial, twice. After each addition, the liquid was swirled around in the vial then added to a clean 250ml separatory funnel. The layers were then allowed to separate. The 80% methanol layer was collected in a new glass scintillation vial. Finally, the methanol layer was allowed to dry under an N^2^ stream completely. After reconstituting the sample in 200μl 100% methanol, the vial was fastened with a lid, secured with Parafilm, labeled, and placed in a −20°c freezer until it was ready for the assay.

Mouse neuroblastoma cells (N2a) (ATCC, CCL131) were cultured and maintained in Eagle’s Minimum Essential Media (EMEM, ATCC) with 10% fetal bovine serum (ATCC) and 5ml penicillin-streptomycin (10,000U/mL) (ThermoFisher Scientific) in a 37°C incubator at 5% CO_2_:95% air atmosphere. The cells were plated at 30,000 cells per well in a 96-well tissue culture plate (Fisher Scientific, 07-200-90). The cells were allowed to incubate overnight in the previously described growth medium. After 18-22 hours of incubation, the cells were treated with either plain medium or medium with Ouabain (31.3μM) and Veratradine (3.13 μM) (O/V), enough to achieve 20% cell death in positive control. Two rows of wells with O/V had the P-CTX3C serial dilution standard added, and four rows of wells (two with O/V and two without O/V) had the extracted samples added. The samples were then incubated overnight.

After 18-22 hours of incubation, the medium was removed from the wells using an electric pump and suction pipette. An MTT (3-[4,5-dimethylthiazole-2-yl]-2,5-diphenyltetrazolium bromide) colorimetric assay was performed, followed by an absorbance reading at 544nm for each well [26]. First, 1ml MTT bromide was added to the 5ml growth medium and then the MTT bromide mixture to each well in 50μl aliquots. The cells were left to incubate for 30-60mins until a purplish color appear. MTT bromide is catalyzed to MTT-formazan by mitochondrial succinate dehydrogenase, which creates a dark purple color. The more metabolically active cells in a well, the darker the color, and therefore the higher the absorbance when measured by a spectrophotometer. After reaching the time limit, the MTT was removed via the flick method and added 100μl of dimethyl sulfoxide (DMSO) to each well. DMSO acts as a lysing agent to the cells that release the color from the cells’ inside. The plate was put on an orbital shaker to distribute the coloring for 15 minutes evenly and read at an absorbance at 544nm.

## Results

### Ciguatera Fish Poisoning Hotspots and Coldspots from Interviews with Informants

In general, the south side of the islands of Puerto Rico, Vieques, and St. Thomas (USVI), and St. John are considered ciguatera fish poisoning (CFP) hotspots by the fishers we interviewed (Figure 1Figure 2). In the USVI, fisheries biologists, Coki Point fishermen, and Frenchtown fishermen all noted that the southern waters of St. Thomas (USVI) were particularly prone to CFP and that the northern waters were not. USVI fisheries biologists stated: “…nearly always, or always, the basic rule is don’t eat the species they listed as toxic in the south but they are all right from the north.” And in Frenchtown, St. Thomas, when asked why the south was toxic, the fishers stated: “…primarily because of the upwellings in the North, which does not allow the ciguatera toxin to grow. Every six months there are upwelling events that kept the toxin away, making the region all right for fish.” One informant from Coki Point, St. Thomas, sorted four fish cards into a pile that comprised those species they do not eat “...if we catch them from the south.” From interviewing three fishermen at Coki Point, the name of a place in the water where fish become toxic was identified as: “Copper Banks” also known as “Scratch and Itch Banks”. When asked about other specific places in the south and they just said, “The whole south.” Yet these informants also said that the eastern waters, east of St. John, were also prone to the toxin. The Frenchtown fishers also said that the toxins grow primarily on the flat surfaces of old wrecks and that this is where a lot of fish feed, but that the currents and occasionally the storms clean the surfaces of the toxin, and a few months later all the fish are good to eat. The fishermen of Frenchtown backed away from saying that it was substrates alone that influenced whether or not the fish they captured might be toxic, suggesting, instead, as is common with local ecological knowledge (LEK), a more advanced and complex understanding of ecosystem dynamics. It is essentially a process that results in ciguatoxin poisoning: a combination of bathymetry, substrates, the behaviors of fish, and the knowledge and behavior of humans (and, perhaps, in some places, their hunger or desperation). Each of these factors combines to result in incidents of human ciguatoxin poisoning. They understand that the toxin ultimately comes from dinoflagellates—they even mentioned these critters by the general name, dinoflagellate—but it isn’t a simple case of fish feeding from them and then passing the toxin on to humans. Instead, it’s the conditions in which the dinoflagellates settle, initially, in that they need the calmer waters of the south and east rather than the seasonal upwellings and currents of the north; storm events can also influence the extent of their distribution, in that often hurricanes “clean” the flat substrates, where they settle and grow, by “tearing up the bottom.”

**Figure 1.**
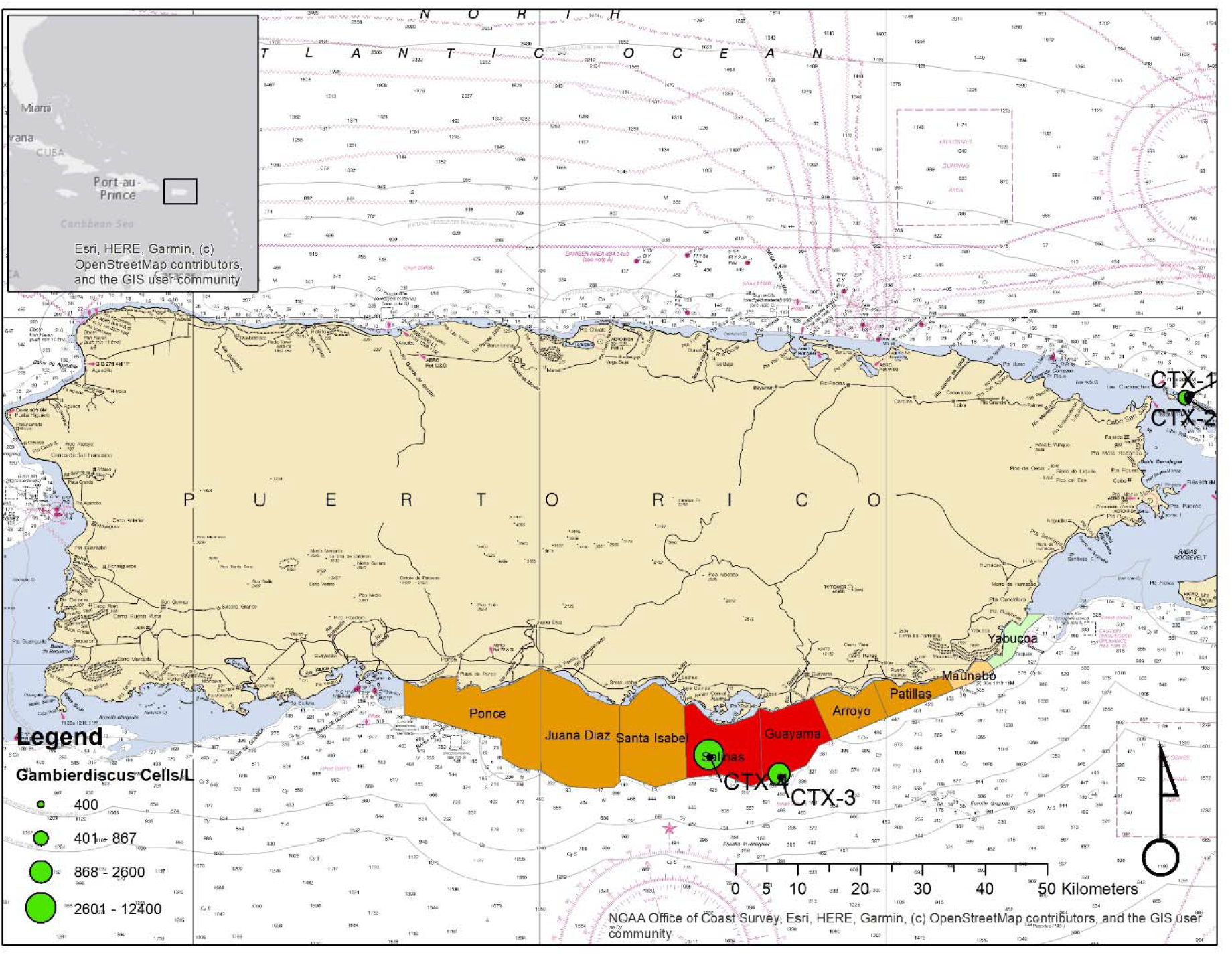
A map of the places named by fishers in our interviews during free-listing of locations with high levels of ciguatoxic fishes. Color of each area shows the liklihood of ciguatoxin (CTX) hotspots as derived from overall salience metrics from fishers’ free-listing of sites with high prevalence of CFP (red 0.34-0.37; orange 0.066-0.095; gold 0.048; yellow 0.034; light green 0.016). Cell counts are shown for all *Gambierdiscus* species on screen rigs deployed at 4 sites: CTX-1 (Fajardo 25m depth), CTX-2 (Fajardo 18m depth), CTX-3 (Guayama 27m depth) and CTX-4 (Salinas 18m depth).

**Figure 2.**
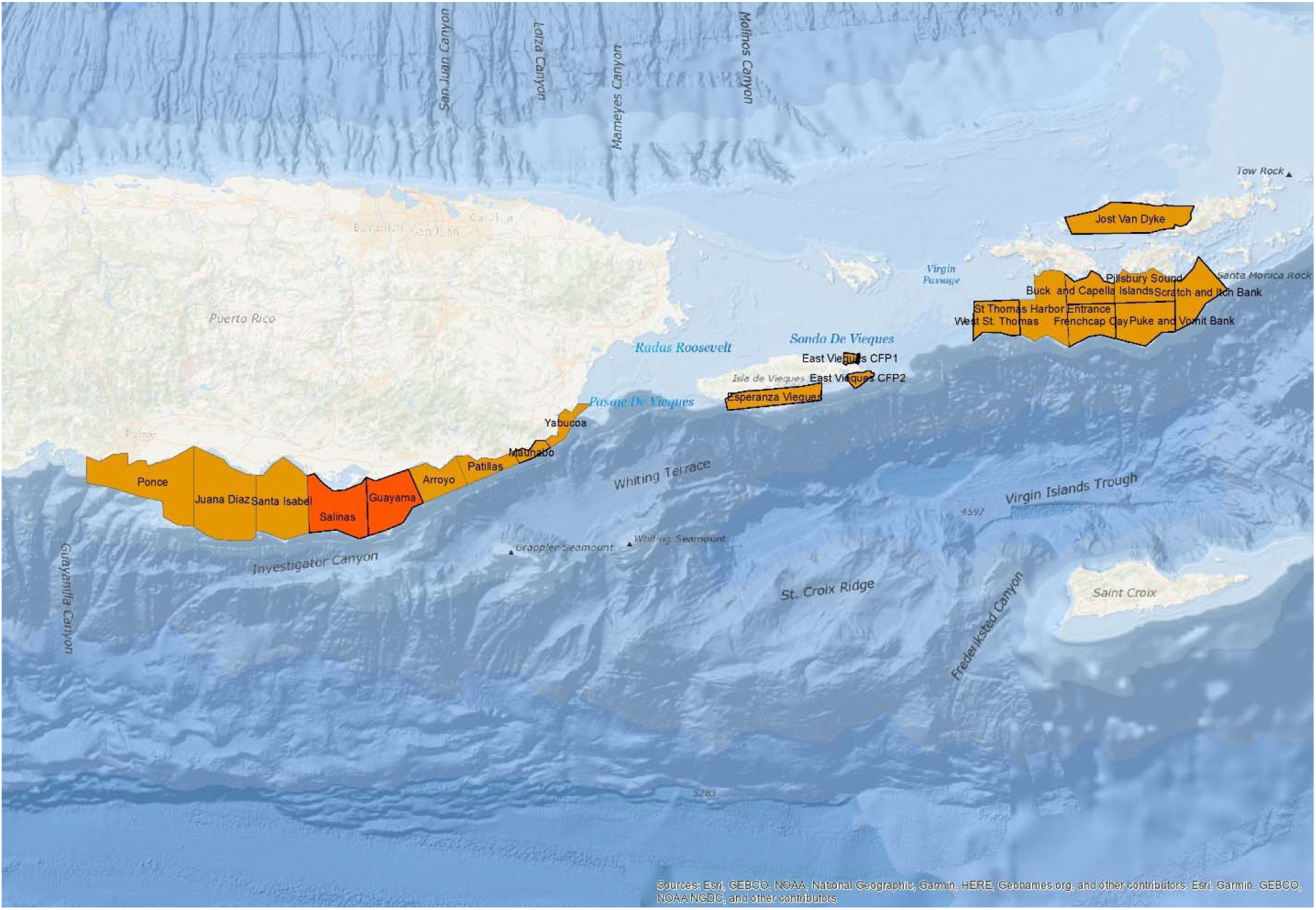
Ciguatera hotspots in Puerto Rico and St. Thomas (USVI) mentioned during interviews with fishers. The red color areas indicate the hotspots which had the highest salience metrics in informant free-lists from Puerto Rico. These hotspots are areas we sampled for CTX in fishes and dinoflagellate abundances. Orange areas were mentioned by informants with less frequency and have not been sampled yet by our team for CTX and dinoflagellates.

The results of the free-listing exercise for hotspot identification show most fishers in Puerto Rico believed Guayama and Salinas were CTX hotspots, and these two locations had the greatest overall salience scores (Table 1). Out of the 21 fishers interviewed, 12 of them identified Guayama as a hotspot area, which was the most frequently mentioned; Guayama had an overall salience score of 0.377. Other areas where CFP has been reported by the fishers are shown in this table along with the salience metric for each location by fishers in different fishing villages used for interview locations. Fishing villages differed in the salience scores associated with these locations, with Guayama fishers rating both Guayama and Salinas highly (salience = 0.53 for Guayama; salience = 0.73 for Salinas), whereas Fajardo fishers rated Vieques as a location with high levels of CFP (salience = 0.62 Table 1). Guayama and Salinas are both on the Southeastern coast of Puerto Rico (Salinas is 15 miles west of Guayama) and fishers from those municipalities share common fishing grounds. Vieques is an island further to the east reachable by boat in 30 min from Fajardo and Naguabo, and fishers from Naguabo travel to those Vieques reefs to fish. In terms of differences among the fishing villages, while fishers on the south coast named Guayama and Salinas as the two most often listed CFP hotspots (salience for Guayama = 0.419, Salinas = 0.386), fishers in the Northeast (Fajardo and Naugabo) named Vieques as a CFP hotspot (salience = 0.546; Table 2). This suggests that there is more than one hotspot in Puerto Rico and the fishing villages experienced a different perception of where hotspots were located, although both agreed that hotspots with high prevalence of CFP were distributed along the south coast of the main island of Puerto Rico and also on the south coast of Vieques (Figure 2). It also means that fishers have a thorough understanding of their fishing grounds. For example, a Fajardo fisher is more knowlegable of Fajardo, Ceiba, Luquillo, Rio Grande, and Culebra fishing grounds. On the other hand, they may have marginal knowledge about fishing grounds in the southern coast and its dynamics. For this study, Guayama was chosen as one hotspot for sampling of dinoflagellates densities, CTX in fish tissues, and ecological food web modeling; however future work needs to be conducted on Vieques and other sites mentioned. Few fishermen chose CFP locations on the eastern, western and northeastern coast of Puerto Rico. No fishermen selected Fajardo as an area of CFP prevalence, which is an area in the Northeast known for its commercial fisheries. For this reason, we chose to investigate reefs to the NE of Fajardo as the coldspot comparison area for ecological sampling.

**Table 1.** Salience metrics for places in Puerto Rico with high levels of ciguatoxic fishes (Locations with CFP) named by fishers during free-listing interviews. Salience is a weighted average of the (inverse) rank of an item across multiple freelists, where each list is weighted by the number of items in the list (Smith 1993). Overall salience (first column) is given for interviews made at all places; salience for individual fish houses or interview locations are listed in the following columns. High salience (in boldface) means good agreement among fishers from a location.

Interview locations
| Locations named<br>in free-lists | Overall | Arroyo | Cabo<br>Rojo | Fajardo | Guayama | Juana<br>Diaz | Maunabo | Naguabo | Ponce |
| --- | --- | --- | --- | --- | --- | --- | --- | --- | --- |
| Guayama | **0.377** | **1** | **0.5** | 0.111 | **0.533** | 0 | **0.667** | **1** | 0 |
| Salinas | **0.337** | 0 | **1** | 0.133 | **0.733** | 0 | 1 | 0 | 0 |
| None | 0.286 | 0 | 0 | 0.2 | 0 | **1** | 0 | 0 | **1** |
| Vieques | 0.156 | 0 | 0 | **0.622** | 0 | 0 | 0 | 0.167 | 0 |
| Ponce | 0.095 | 0 | 0 | 0.2 | 0.167 | 0 | 0 | 0 | 0 |
| Juana Diaz | 0.08 | 0 | 0 | 0.178 | 0.133 | 0 | 0 | 0 | 0 |
| Santa Isabel | 0.066 | 0 | 0 | 0.156 | 0.1 | 0 | 0 | 0 | 0 |
| Arroyo | 0.056 | 0 | 0 | 0 | 0 | 0 | 0.333 | **0.833** | 0 |
| Patillas | 0.048 | 0 | 0 | 0.067 | 0 | 0 | 0 | **0.667** | 0 |
| Maunabo | 0.034 | 0 | 0 | 0.044 | 0 | 0 | 0 | 0.5 | 0 |
| Arroy | 0.021 | 0 | 0 | 0.089 | 0 | 0 | 0 | 0 | 0 |
| Yabucoa | 0.016 | 0 | 0 | 0 | 0 | 0 | 0 | 0.333 | 0 |

**Table 2.** Salience metrics for fishers interviewed along the North and South coasts in Puerto Rico who were asked to name places with high levels of ciguatoxic fishes during interviews. Overall salience (first column) is given for interviews made at all places; salience metrics from coast-wide pooled interviews (North Coast and South Coast) are listed in the other columns.

| Item | OVERALL | North Coast | South Coast |
| --- | --- | --- | --- |
| Guayama | **0.377** | 0.259 | **0.424** |
| Salinas | **0.337** | 0.111 | **0.427** |
| None | 0.286 | 0.167 | **0.333** |
| Vieques | 0.156 | **0.546** | 0 |
| Ponce | 0.095 | 0.167 | 0.067 |
| Juana Diaz | 0.08 | 0.148 | 0.053 |
| Santa Isabel | 0.066 | 0.13 | 0.04 |
| Arroyo | 0.056 | 0.139 | 0.022 |
| Patillas | 0.048 | 0.167 | 0 |
| Maunabo | 0.034 | 0.12 | 0 |
| Arroy | 0.021 | 0.074 | 0 |
| Yabucoa | 0.016 | 0.056 | 0 |

The consensus analysis results on the fish card pile-sort exercise indicated a strong agreement among the fishers on the fish species most likely to be ciguatoxic (**Error! Reference source not found.**). The large eigenratio (18.35) and the lack of negative competence scores (Table 3) in the consensus analysis indicate a good fit for the consensus model. The fish species that informants agreed upon as high prevalence for CFP were hogfish (*Lachnolaimus maximus*), barracuda (*Sphyraena barracuda*), king mackerel (*Scomberomorus cavalla*), black jack (*Caranx lugubris*), greater amberjack (*Seriola dumerili*), and horse-eye jack (*Caranx latus*); these were the species in the “answer key” generated by the consensus analysis as being toxic (Table 4). All of these fish species plotted in the center of the consensus analysis network, indicating that informants from all fishing villages thought that those species were toxic. These also had very high salience scores on the free-listing analysis, with great barracuda having the highest salience score (0.898) followed by hogfish (0.728) and amberjack (0.488; Table 4). There were several species named in free-lists that were not on our pile sort cards: African pomano, bar jack, yellow goatfish, almaco jack, escolar, and rainbow runner, but these all had relatively low salience scores. There were some differences noted amongst the fishers in the different villages; fishers from Guayama tended to rate schoolmaster (salience = 0.081), king mackerel (salience = 0.215), and dog snapper (salience = 0.179) as toxic fishes, while fishers from Fajardo identified yellowfin grouper and rainbow parrotfish (neither were listed on free-lists) as toxic. A fisher from Maunabo identified Cubera snapper and tiger grouper (neither were listed on free-lists) as toxic. Fishers from Ponce and Maunabo also identified king mackerel and dog snapper as toxic, agreeing with the Guayama fishers. Fishers from Juana Diaz and Naguabo identified cero as toxic (salience = 0.018).

**Table 3.** Consensus analysis results: Informant ID, fish house location, years of commercial fishing experience and competence scores for the informants interviewed about CFP in fishes caught in Puerto Rico.

| Informant ID | Fish house | Years of<br>Experience | Competence Score |
| --- | --- | --- | --- |
| 1 | CaboRojo | 47 | 0.97 |
| 2 | Fajardo | 39 | 0.854 |
| 3 | Fajardo | 67 | 0.97 |
| 4 | Fajardo | 33 | 0.883 |
| 5 | Fajardo | 36 | 0.924 |
| 6 | Fajardo | 40 | 0.924 |
| 7 | Guayama | 18 | 0.93 |
| 8 | Guayama | 28 | 0.926 |
| 9 | Guayama | 32 | 0.93 |
| 10 | Guayama | 25 | 0.93 |
| 11 | Guayama | 23 | 0.93 |
| 12 | Guayama | 30 | 0.974 |
| 13 | Arroyo | 58 | 0.974 |
| 14 | Arroyo | 27 | 0.933 |
| 15 | JuanaDiaz | 44 | 0.855 |
| 16 | JuanaDiaz | 31 | 0.97 |
| 17 | JuanaDiaz | 25 | 0.868 |
| 18 | JuanaDiaz | 45 | 0.952 |
| 19 | Ponce | 37 | 0.974 |
| 20 | Maunabo | 20 | 0.836 |
| 21 | Naguabo | 24 | 0.855 |
|  | Average | 34.7 | 0.922 |

**Table 4.** Comparison of fishers’ perception of which fish species are ciguatoxic based on pile-sort exercise and free-listing exercise (salience metric).

|  | **Pile-sort Species** | **Answer**<br>**key** | **Free-list Species** | **Salience** |
| --- | --- | --- | --- | --- |
| 1 | Hogfish | 1 | Hogfish | 0.728 |
| 2 | Barracuda | 1 | Barracuda | 0.898 |
| 3 | King Mackerel | 1 | King Mackerel | 0.215 |
| 4 | Cero | 0 | Cero | 0.018 |
| 5 | Black Jack | 1 | Black Jack | 0.243 |
| 6 | Amberjack | 1 | Amberjack | 0.488 |
| 7 | Bluerunner | 0 |  |  |
| 8 | Horse-eye Jack | 1 | Horse-eye Jack | 0.143 |
| 9 | Jack Crevalle | 0 |  |  |
| 10 | Cubera Snapper | 0 |  |  |
| 11 | Queen Snapper | 0 |  |  |
| 12 | Silk Snapper | 0 |  |  |
| 13 | Blackfin Snapper | 0 |  |  |
| 14 | Lane Snapper | 0 |  |  |
| 15 | Mutton Snapper | 0 |  |  |
| 16 | Mangrove Snapper | 0 |  |  |
| 17 | Yellowtail Snapper | 0 |  |  |
| 18 | Schoolmaster | 0 | Schoolmaster | 0.081 |
| 19 | Dog Snapper | 0 | Dog Snapper | 0.179 |
| 20 | Tiger Grouper | 0 | **Species not on cards** |  |
| 21 | Red Hind | 0 | African Pompano | 0.110 |
| 22 | Coney | 0 | Cobia (Rainbowrunner) | 0.015 |
| 23 | Yellowfin Grouper | 0 | Bar Jack | 0.041 |
| 24 | Queen Parrotfish | 0 | Yellow Goatfish | 0.032 |
| 25 | Rainbow Parrotfish | 0 | Almaco Jack | 0.024 |
| 26 | Stoplight Parrotfish | 0 | Escolar | 0.024 |
| 27 | Striped Mojarra | 0 |  |  |
| 28 | Yellowfin Mojarra | 0 |  |  |
| 29 | Sand Tilefish | 0 |  |  |
| 30 | Spadefish | 0 |  |  |
| 31 | Trunkfish | 0 |  |  |
| 32 | Redear Sardine | 0 |  |  |
| 33 | White Mullet | 0 |  |  |
| 34 | Ballyhoo | 0 |  |  |
| 35 | Blue Crab | 0 |  |  |
| 36 | Queen Conch | 0 |  |  |
| 37 | West Indian Topshell | 0 |  |  |

The number of *Gambierdiscus spp.* cells L^−1^in the hotspots were higher than in the coldspots (Figure 4). The median values in coldspot CTX-1 and coldspot CTX-2 were 333.33 cells L^−1^ and 1000 cells L^−1^, respectively. The hotspots’ median values were higher at 2333.33 cells L^−1^ at CTX-3 and 11,666.67 cells L^−1^ at CTX-4. The short boxes in sites CTX-1, CTX-2, and CTX-3 show a high agreement among the replicate samples, while CTX-4 suggests more considerable differences in the repeats. The lower whisker in the CTX-3 plot site overlaps the first quartile in the CTX-2 site plot. These data show that there are some similar cell counts in CTX-3 and CTX-2. The CTX-4 site had many more cells L^−1^ than any other site.

The estimated CTX levels in the targeted species *S. barracuda* were higher in the hotspot than in the coldspot by .071 CTX-3C equiv. (Welch Two Sample t-test, n = 8, p = 0.03834) (Figure 5). Median values of *S. barracuda, Caranx ruber, and L. maximus* were all higher in the hotspot than the same species in the coldspot. *Holocentrus rufus* and *Sparisoma viride* did not differ between sites. A two-way interaction ANOVA showed a significant effect of hotspot/coldspot on toxin concentration in fishes (F = 6.359, f = 1. P = 0.016) as well as an effect of trophic group on toxin concentration in fishes (F = 5.078, df = 2, p = 0.0111).

**Figure 3.**
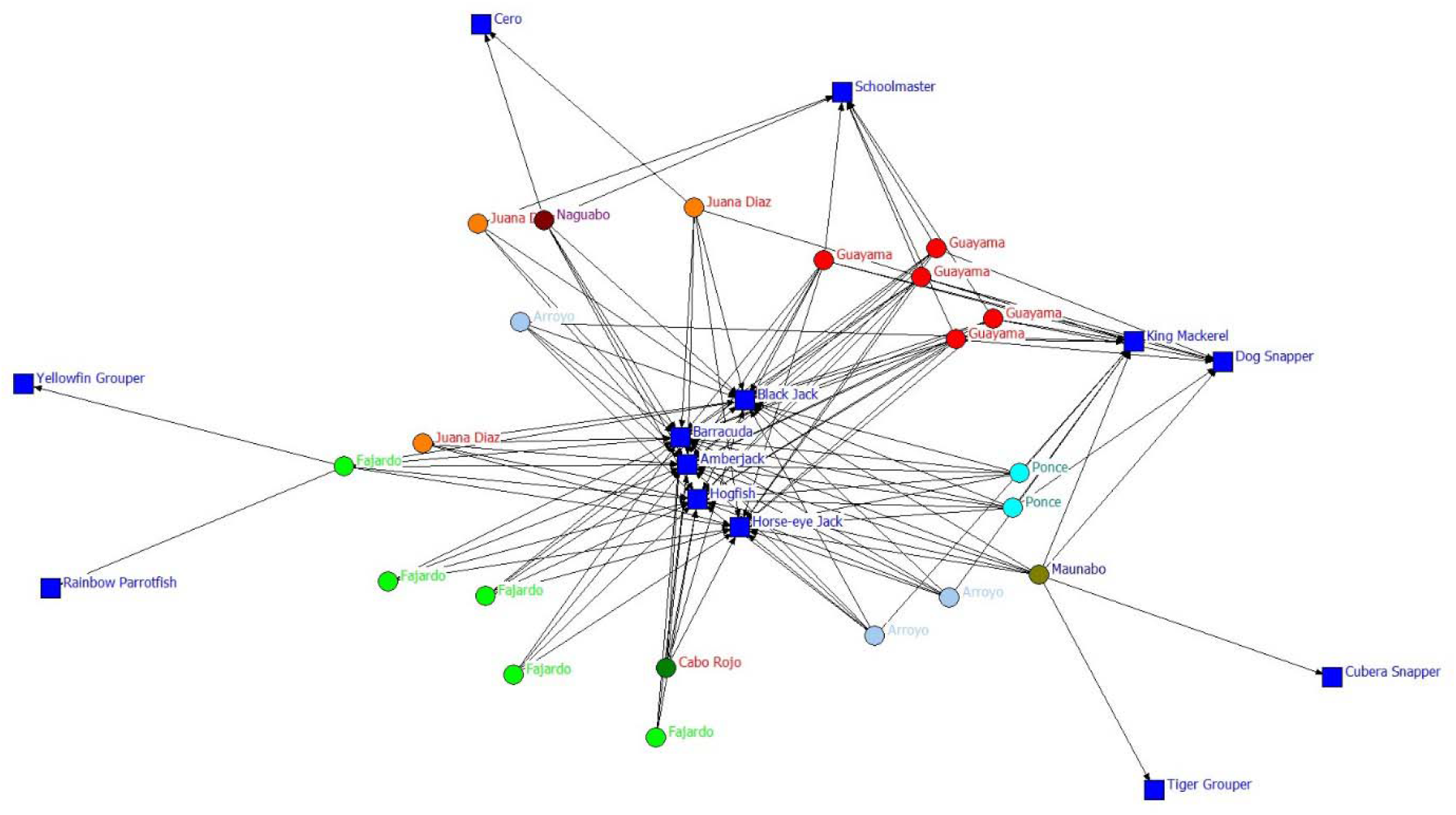
The consensus analysis of fish species identified by informants as having a high prevalence of CFP in the pile-sort exercise. Fish species were offered as photo cards to be sorted into toxic and non-toxic piles and are shown as blue squares. The colored circles represent individual informants and are labled and color-coded by their fishing villages. Lines show which fishers linked a species to CFP. Many lines linking a fish species to the informants indicates a consensus among those fishers that the fish species is toxic. A greater consensus is found among informants for CFP in fish species at the center of this plot than at the edges of the plot, which were species infrequently identified as having CFP.

**Figure 4.**
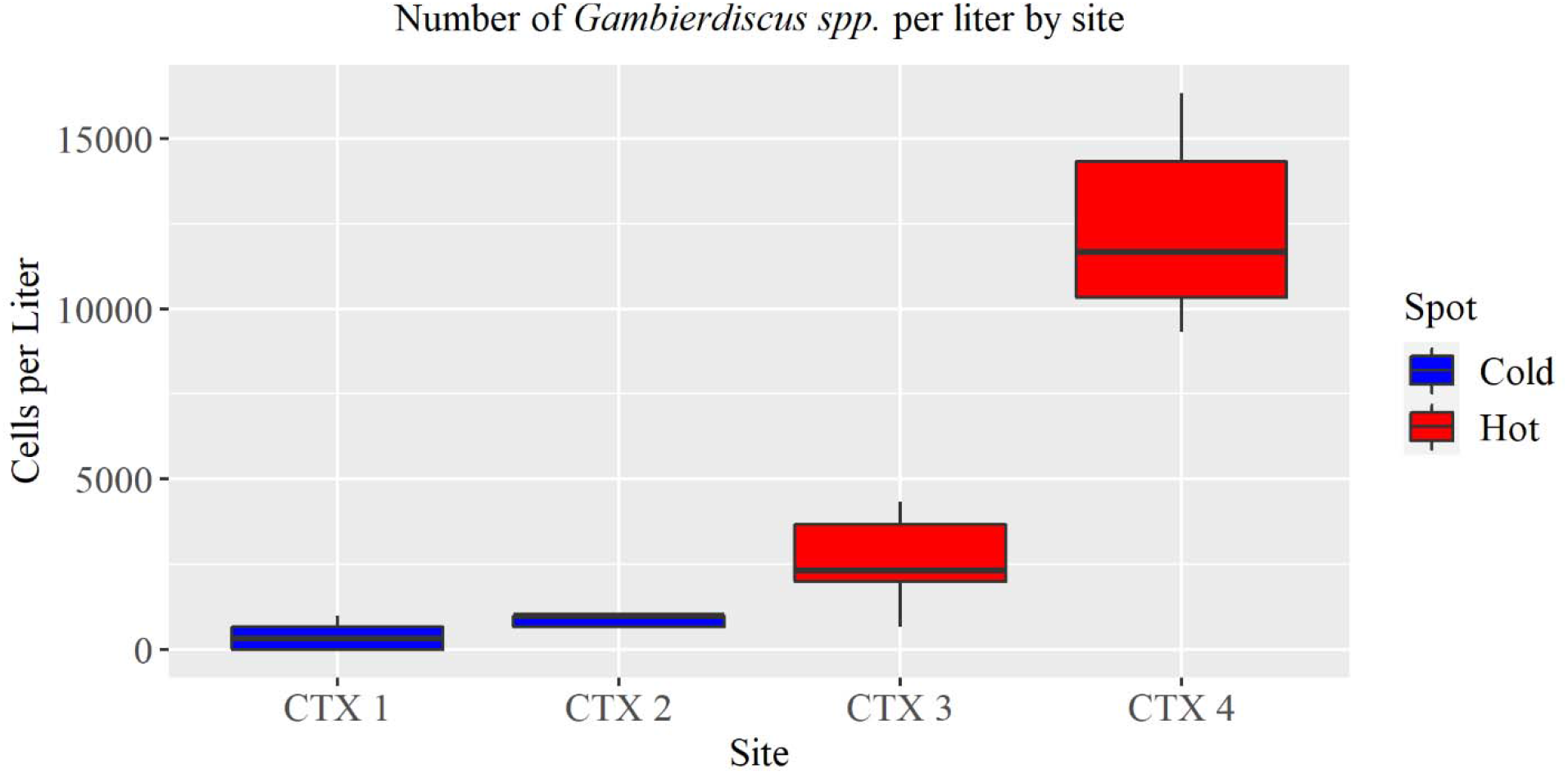
Results for the dinoflagellate cell count differences between hotspot and coldspot. *Gambierdiscus* spp. cells L^−1^for the coldspots (CTX-1 and CTX-2) and the hotspots (CTX-3 and CTX-4). The experts at the NOAA Southeast Fisheries Laboratory (Beaufort, NC) counted the cells and confirmed the cells are in the *Gambierdiscus* genera.

**Figure 5.**
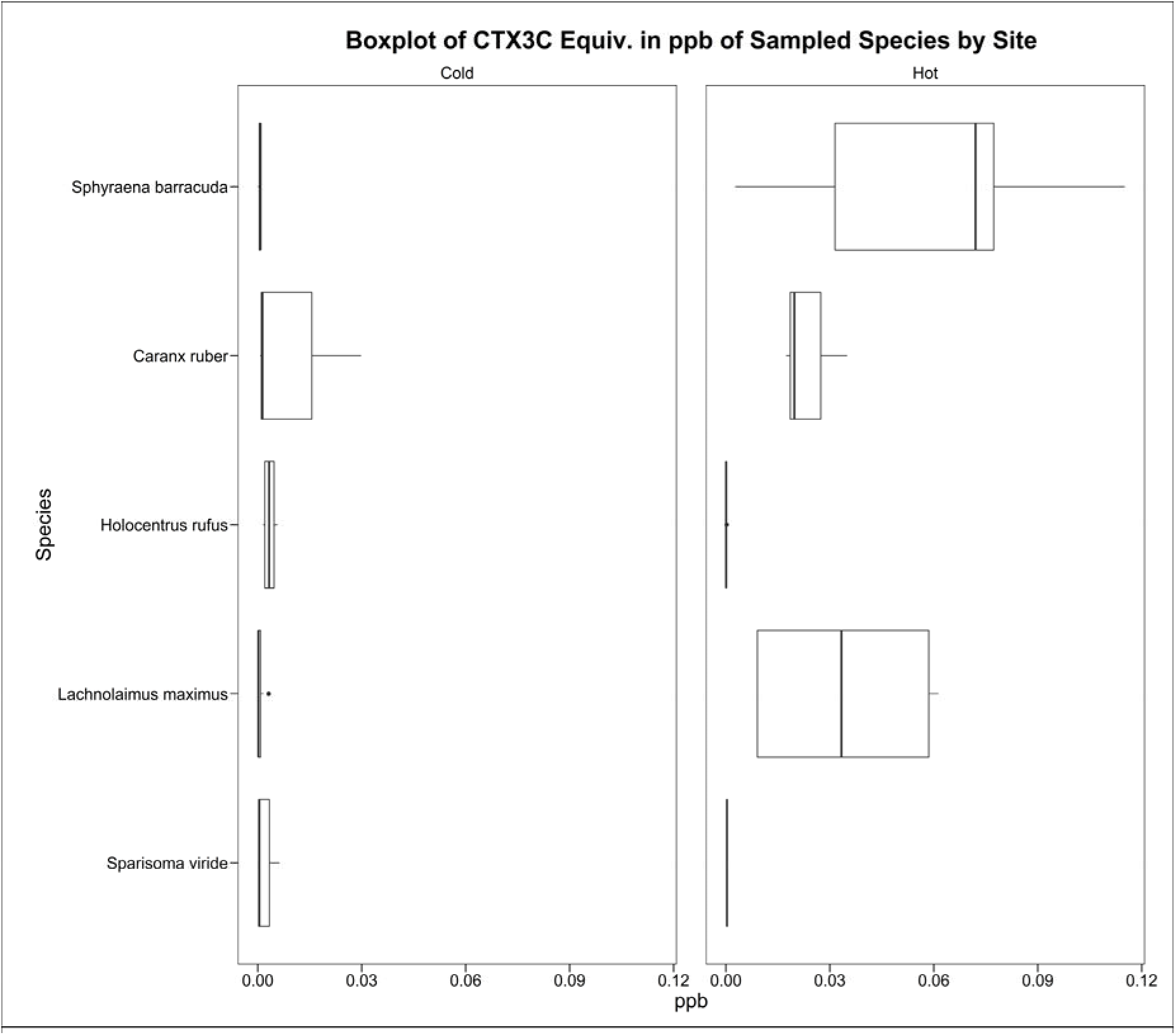
Boxplot of median CTX3C equiv. concentrations in ppb by species in the hotspot and coldspot. The top trophic predators had a higher median CTX3C equiv. concentration in the hotspot compared to the coldspot. Species are listed from highest ETL to lowest.

As the trophic level increases, the CTX concentration increases (Figure 6). The estimated CTX values in the coldspot did not differ as trophic level increased. However, a Tukey HSD post-hoc comparison showed that it did in the hotspot. The low and medium trophic groups in the hotspot both had lower toxin concentrations than the high trophic group in the hotspot (p = 0.029 and p = 0.027, respectively).

**Figure 6.**
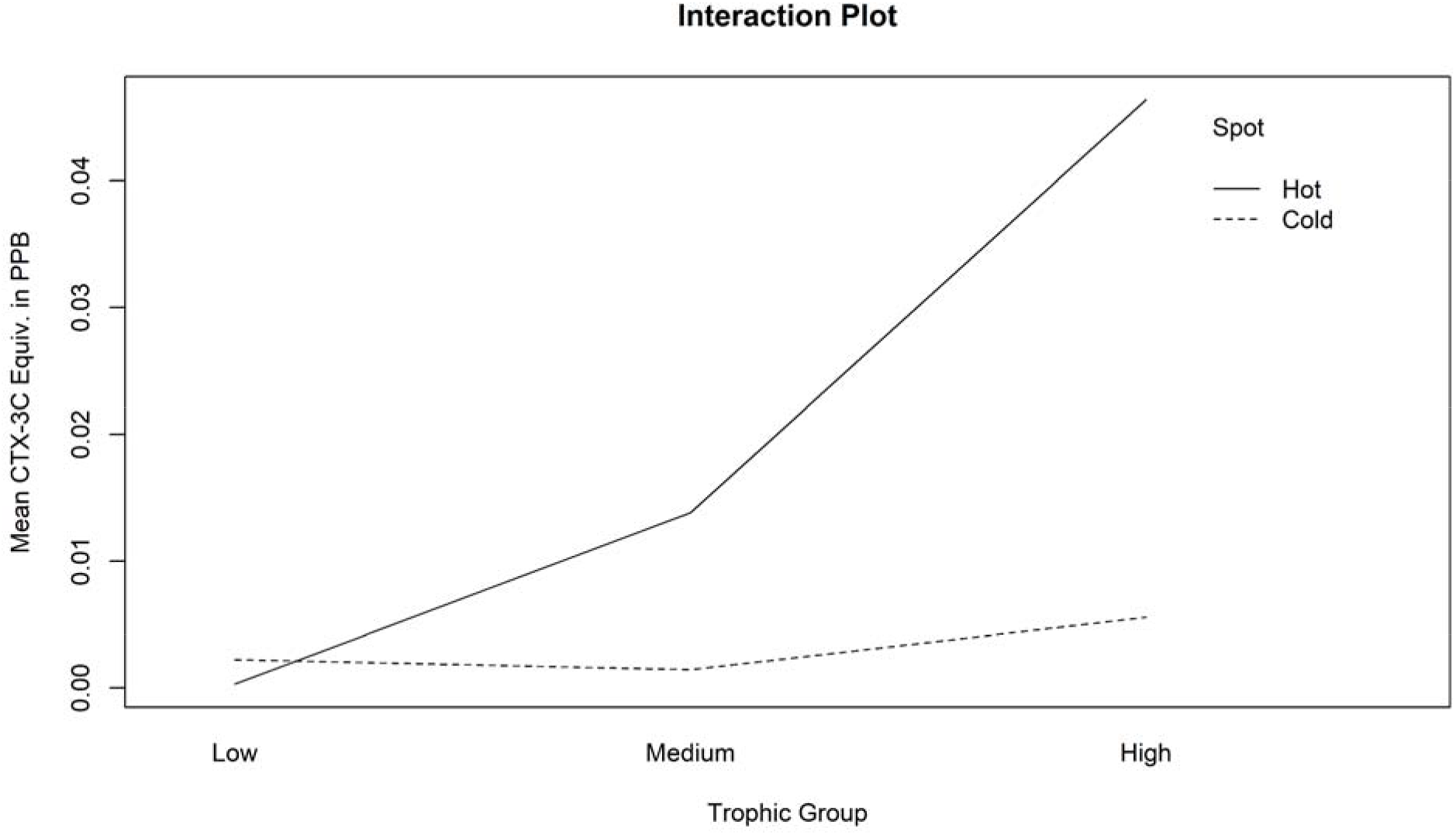
Interaction plot of CTX3C equiv. concentrations in three trophic level groups (low, medium, and high) between the hotspot and coldspot.

## Discussion

### Fishers know where CTX is more prevalent

Fishers along the east coast of Puerto Rico identified Guayama as a CTX hotspot (Fig 1) including fishers from Cabo Rojo to Guayama and the northeast coast of Fajardo. There haven’t been formal ciguatera hotspot identification studies done in Puerto Rico or any other Caribbean islands, only in the Pacific in Hawaii. More research is done on ciguatoxin in the Pacific than in the Caribbean due to more funding allocation, and P-CTX’s are 10-fold more toxic than C-CTX’s [27]. There are also readily available Pacific ciguatoxin standards available for purchase to run assays with (P-CTX-3C, Fujifilm Wako Chemicals), but none is available for Caribbean chemical strains. The lack of data emphasizes the importance of fishers’ LEK, which can improve fisheries’ management and can be used in data-poor artisanal fisheries to understand fishing grounds [28,29].

Fishers across Puerto Rico generally agreed on which locations were more likely to be toxic and which fishes to avoid, which means there is some form of data transmission. Information could be passed through fish houses to fishers at other fish houses, rumors of locals getting sick and tracing that illness back to which fish house the specimen was purchased from, or passed down from elders in the communities. We argue that LEK and TEK provides a community adaptation to CFP that leads to a reduction in the number of CFP cases, although insufficient data on CFP cases make this challenging to investigate. Our field work made clear that most fishers avoid catching “hot fish” at “hotspots” which reduces the incidence rates of CFP. Casual consumers do not know as much about CTX as commercial fishers. Researchers should investigate the knowledge gap between commercial and recreational fishers. Non-commercial fishers with less knowledge may keep riskier fishes leading to CFP outbreaks.

It is difficult to determine which fish are toxic when there are no dockside tests available, potentially leading to a deterministic view of ciguatera fish poisoning [30]. Nellis and Bernard (1986) [30] show that in the USVI, when it comes to CFP, people believed they would eventually get it, and there wasn’t much they could do about it.. There was a similar sentiment from fishers in Puerto Rico. Their methods of avoiding toxic fish could only go so far; catching a contaminated fish was inevitable. One informant in Guayama mentioned that he had CFP multiple times. In addition to selling his catch, he also fed his family with the fish he caught. To prevent them from getting sick he would eat the fish first, freeze the rest, then wait a few days before feeding it to his family, to make sure they wouldn’t get sick. However, his fishing style didn’t change, partially because he couldn’t change it. Those who live in the more impoverished areas cannot change their fishing locations due to economic restraints. They generally fish from smaller boats and do not have the means to trailer it to safer fishing gounds. They rely on knowledge they have gained as well as knowledge that has been passed down to them as a way to avoid catching toxic fish.

### Between site dinoflagellate counts

Overall, the data shows higher cell counts of low toxicity species in the hotspot samples than in the coldspot samples. The higher number of cells L^−1^ could be causing toxicity in higher trophic level fishes at those sites. Herbivores and herbivorous fish consume these dinoflagellates when feeding on their preferred substrates, which means any increase in the number of cells resting on the algae would increase the amount of toxin entering the system [31–33].

The suite of species found was different at each site, with *G. caribaeus* being the only species found at both hotspot and coldspot (Figure 7). Litaker *et al*. 2017 describe each species’ toxin concentration that we found at the hotspot and coldspot: the toxin concentration of *Gambierdiscus caribaeus* is 0.66 ± 0.34 fg CTX3C equiv. cell^−1^, *Gambierdiscus carpenteri* is 0.89 ± 0.41 fg CTX3C equiv. cell^−1^ *Gambierdiscus belizeanus* is 0.85 ± 0.81 fg CTX3C equiv. cell^−1^ and *Gambierdiscus carolinianus* is 0.27 ± 0.43 fg CTX3C equiv. cell^−1^. Assuming an equal distribution of cells, although unlikely, the average toxicity of the cells at the coldspot (0.8 fg CTX3C equiv. cell^−1^) is almost twice as high as the cell toxicity at the hotspot (0.465 fg CTX3C equiv. cell^−1^). Since this is counterintuitive to what we predicted, there may be more toxic species cells than low toxic species in the hotspot. Future studies should include more in-depth dinoflagellate sampling protocols, including doing the qPCR right after the cells are captured (our qPCR was delayed due to the global SARS-CoV-2 pandemic, and therefore, some DNA was degraded).

**Figure 7.**
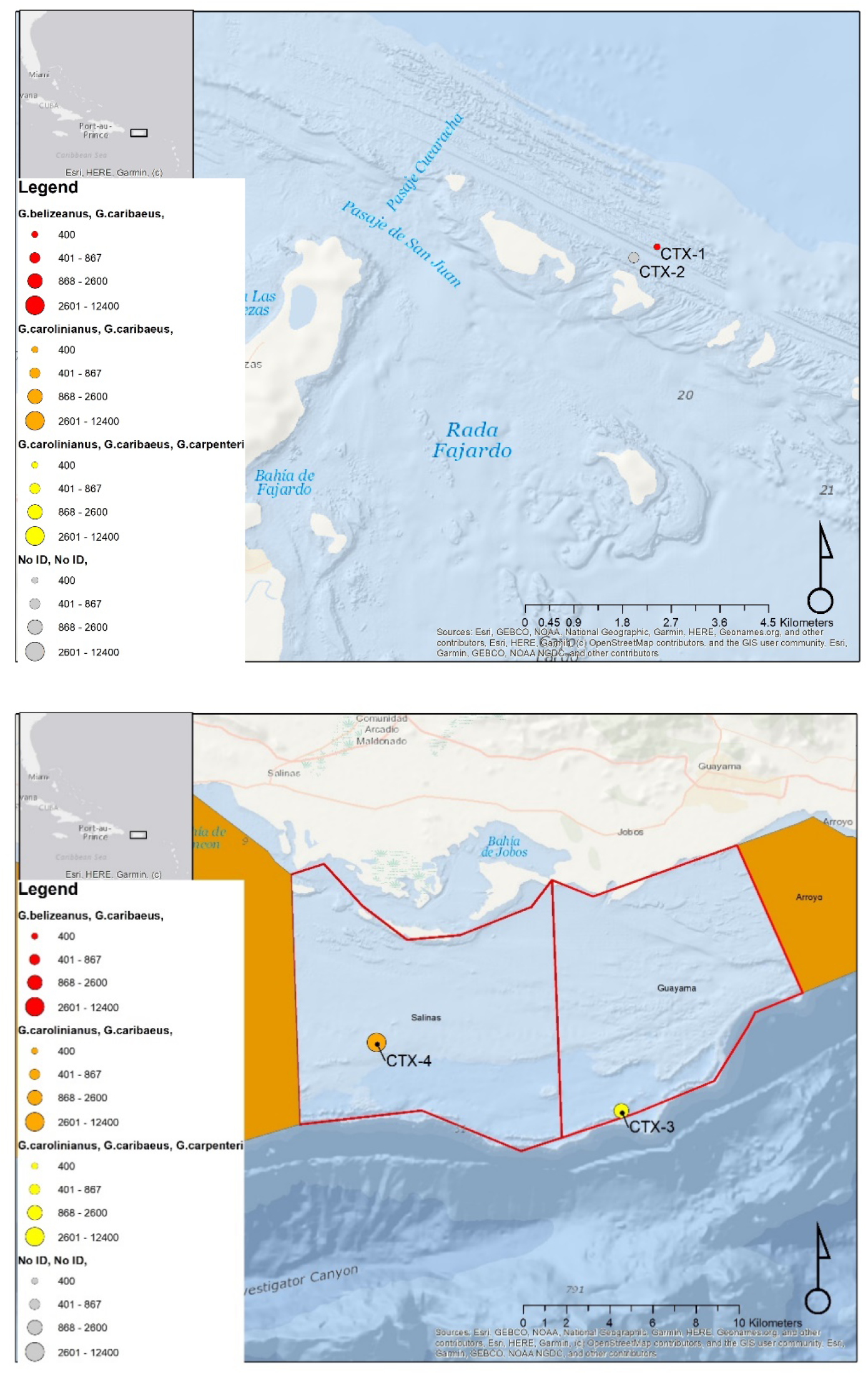
Map of the screen rig sites and the Gambierdiscus spp. identified at those sites. Top: Coldspot sites; CTX-1 was 25m deep, CTX-2 was 18 m deep. Bottom: Hotspot sites; CTX-3 was 27 m deep and CTX-4 was 18 m deep. CTX-2 species were not identified by qPCR due to the samples’ degraded DNA.

If the number of cells of these toxin-producing dinoflagellates drives the toxicity in high trophic level fishes, scientists will benefit from a routine monitoring program of the algae. Divers should collect *Gambierdiscus* spp. using the screen-sampler method, count the number of cells, and identify the species present using PCR, which would also help fill the large data gap in these cells’ global distribution [23]. We generally know which species habituate the Pacific, Atlantic, and Indian Oceans and the Caribbean Sea. However, scientists know little about the specific reefs and coasts to which these dinoflagellates thrive. There is some evidence that increased wave and wind action reduces the toxicity of reefs [34]; northern coasts of the Caribbean Islands experience harsher conditions, disturbing the growth of these algae. Studies should sample along the north and south coasts of Puerto Rico and compare the dinoflagellate profiles to the wind and wave energy exerted on these areas.

### Toxicity in fishes

Overall, fishes in the hotspot had higher levels of CTX in their tissues than the fishes in the coldspot, which supports our hypothesis that fishers can identify CTX hotspots and coldspots. It is difficult to pinpoint what factors drive this difference, but some could be attributed to the higher cell densities in the hotspot than the coldspot. The coldspot in Puerto Rico is on the north side of the island, and a study by Loeffler *et al*. (2018) [34] shows lower toxicity in fishes collected on the north side of the U.S. Virgin Islands. The scientists in this study show greater wave energy on the north side of the USVI, which could be leading to a more deficient growing environment for CTX-producing dinoflagellates.

The fishes that differed from the hotspot and coldspot were the *Sphyraena barracuda* (barracuda), *Lachnolaimus maximus* (hogfish), and *Caranx ruber* (bar jack). These species are all higher trophic level organisms compared to the other species compared. The barracuda consumes mostly fishes with some octopuses and crustaceans, similar to the bar jack, while the hogfish primarily consume mollusks [35]. Interestingly, hogfish have higher levels of CTX3C equiv. in their tissues than the bar jack when the bar jack is at a higher trophic level. When secondary consumers feed on the CTX-producing dinoflagellates, they metabolize the toxin and excrete 95% in the form of oxocenes, which drastically reduces the amount of CTX that gets transferred to the next trophic levels [33]. However, the same metabolism is most likely not present in gastropods, the hogfishes’ preferred prey. Suppose gastropods consume toxin-producing dinoflagellates while grazing on their preferred substrates and are not metabolizing it like fishes. In that case, they could be transferring more CTX to higher trophic levels than if it had gone through herbivorous fishes. This CTX transfer could explain the higher levels of CTX3C equiv. in hogfish compared to the bar jack. Future studies should test the CTX3C equiv. concentration in gastropods and secondary consumers in the same locations and compare that to the dinoflagellate density and species composition on the same reef. This study may begin to explain how the pathways that CTX takes through the food web play a role in the toxicity of some species.

## Notes

https://osf.io/btyd3/

### Competing Interest Statement

The authors have declared no competing interest.

## References

1. Lewis RJ, Sellin M, Poli MA, Norton RS, MacLeod JK, Sheil MM. Purification and Characterization of Ciguatoxins from Moray Eel (Lycodontis javanicus, Muraenidae). Toxicon. 1991;29: 1115–1127. doi:10.1016/0041-0101(91)90209-A

2. Pottier I, Vernoux JP, Jones A, Lewis RJ. Characterisation of Multiple Caribbean Ciguatoxins and Congeners in Individual Specimens of Horse-Eye Jack (Caranx latus) by High-Performance Liquid Chromatography/Mass Spectrometry. Toxicon. 2002;40: 929–939. doi:10.1016/S0041-0101(02)00088-0

3. Lehane L, Lewis RJ. Ciguatera: Recent Advances but the Risk Remains. Int J Food Microbiol. 2000;61: 91–125. doi:10.1016/S0168-1605(00)00382-2

4. de Fouw JC, van Egmond HP, Speijers GJA. Ciguatera Fish Poisoning: a Review. RIVM Rep. 2001; 1–66. Available: http://rivm.openrepository.com/rivm/handle/10029/9457

5. Murata M, Legrand a. M, Ishibashi Y, Fukui M, Yasumoto T. Structures and Configurations of Ciguatoxin from the Moray Eel Gymnothorax javanicus and its likely Precursor from the Dinoflagellate Gambierdiscus toxicus. J Am Chem Soc. 1990;112: 4380–4386. doi:10.1021/ja00167a040

6. Lewis RJ. Structure of Caribbean Ciguatoxin Isolated from Caranx latus. J Am Chem Soc. 1998;120: 5914–5920. doi:10.1021/ja980389e

7. Lewis RJ. Ciguatera Management. SPC Live Reef Fish Inf Bull. 2000;7.

8. Copeland NK, Palmer WR, Bienfang PK. Ciguatera Fish Poisoning in Hawai’i and the Pacific. Hawaii J Med Public Health. 2014;73: 24–27.

9. Darius HT, Drescher O, Ponton D, Pawlowiez R, Laurent D, Dewailly E, et al. Use of Folk Tests to Detect Ciguateric Fish: a Scientific Evaluation of their Effectiveness in Raivavae Island (Australes, French Polynesia). Food Addit Contam Part A. 2013;0049: 37–41. doi:10.1080/19440049.2012.752581

10. Hardison DR, Holland WC, McCall JR, Bourdelais AJ, Baden DG, Darius HT, et al. Fluorescent Receptor Binding Assay for Detecting Ciguatoxins in Fish. PLoS One. 2016;11: 1–19. doi:10.1371/journal.pone.0153348

11. Pawlowiez R, Darius HT, Cruchet P, Rossi F, Caillaud A, Laurent D, et al. Evaluation of Seafood Toxicity in the Australes Archipelago (French Polynesia) Using the Neuroblastoma Cell-based Assay. Food Addit Contam - Part A Chem Anal Control Expo Risk Assess. 2013;30: 567–586. doi:10.1080/19440049.2012.755644

12. Reverté L, Soliño L, Carnicer O, Diogène J, Campàs M. Alternative Methods for the Detection of Emerging Marine Toxins: Biosensors, Biochemical Assays and Cell-Based Assays. Marine Drugs. 2014. doi:10.3390/md12125719

13. Davis A, Wagner JR. Who Knows? On the Importance of Identifying “Experts” When Researching Local Ecological Knowledge. Hum Ecol. 2003;31: 463–489.

14. Charnley S, Fischer AP, Jones ET. Integrating Traditional and Local Ecological Knowledge into Forest Biodiversity Conservation in the Pacific Northwest. For Ecol Manage. 2007;246: 14–28. doi:10.1016/j.foreco.2007.03.047

15. Garcia-Quijano CG. Fishers’ Knowledge of Marine Species Assemblages: Bridging between Scientific and Local Ecological Knowledge in Southeastern Perto Rico. Am Anthropol. 2007;109: 529–536. doi:10.1525/AA.2007.109.3.529.530

16. Bergmann M, Hinz H, Blyth RE, Kaiser MJ, Rogers SI, Armstrong M. Using Knowledge from Fishers and Fisheries Scientists to Identify Possible Groundfish Essential Fish Habitats. Fish Res. 2004;66: 373–379. doi:10.1016/j.fishres.2003.07.007

17. Rasalato E, Maginnity V. Using Local Ecological Knowledge to Identify Shark River Habitats in Fiji (South Pacific). Environ Conserv. 2019;37: 90–97. doi:10.1017/S0376892910000317

18. Borgatti S. ANTHROPAC 4.0. Analytic Technologies. Natick, MA; 1996.

19. Smith JJ. Using ANTHROPAC 3.5 and a spreadsheet to compute a freelist salience index. Cult Anthropol Methodol Newsl. 1993;5: 1–3.

20. Borgatti SP, Everett MG, Freeman LC. Ucinet for Windows: Software for social network analysis. Lexington, KY, USA: Analytic Technologies; 2002.

21. Borgatti SP. NetDraw: Graph Visualization Software. Harvard, MA: Analytic Technologies; 2002.

22. Tester PA, Kibler SR, Holland WC, Usup G, Vandersea MW, Leaw CP, et al. Sampling harmful benthic dinoflagellates: Comparison of artificial and natural substrate methods. Harmful Algae. 2014;39: 8–25. doi:10.1016/j.hal.2014.06.009

23. Litaker RW, Vandersea MW, Faust MA, Kibler SR, Nau AW, Holland WC, et al. Global Distribution of Ciguatera Causing Dinoflagellates in the Genus Gambierdiscus. Toxicon. 2010;56: 711–730. doi:10.1016/j.toxicon.2010.05.017

24. Vandersea MW, Kibler SR, Holland WC, Tester PA, Schultz TF, Faust MA, et al. Development of semi-quantitative pcr assays for the detection and enumeration of gambierdiscus species (gonyaulacales, dinophyceae). J Phycol. 2012;48: 902–915. doi:10.1111/j.1529-8817.2012.01146.x

25. Opitz S. Trophic Interactions in Caribbean Coral Reefs Trophic Interactions in Caribbean Coral Reefs. 1996.

26. Pisapia F, Holland WC, Hardison DR, Litaker RW, Fraga S, Nishimura T, et al. Toxicity Screening of 13 Gambierdiscus Strains Using Neuro-2a and Erythrocyte Lysis Bioassays. Harmful Algae. 2017;63: 173–183. doi:10.1016/j.hal.2017.02.005

27. Lewis RJ, Jones A, Vernoux JP. HPLC/Tandem Electrospray Mass Spectrometry for the Determination of Sub-PPB Levels of Pacific and Caribbean Ciguatoxins in Crude Extracts of Fish. Anal Chem. 1999;71: 247–50. doi:10.1021/ac980598h

28. Silvano RAM, Valbo-Jørgensen J. Beyond fishermen’s tales: Contributions of fishers’ local ecological knowledge to fish ecology and fisheries management. Environ Dev Sustain. 2008;10: 657–675. doi:10.1007/s10668-008-9149-0

29. Pita P, Fernández-Vidal D, García-Galdo J, Muíño R. The Use of the Traditional Ecological Knowledge of Fishermen, Cost-Effective Tools and Participatory Models in Artisanal Fisheries: Towards the Co-management of Common Octopus in Galicia (NW Spain). Fish Res. 2016;178: 4–12. doi:10.1016/j.fishres.2015.07.021

30. Nellis DW, Barnard GW. Ciguatera: A Legal and Social Overview. Marine Fisheries Review. 1986. pp. 2–5.

31. Randall JE. A Review of Ciguatera, Tropical Fish Poisoning, with a Tentative Explanation of its Cause. Bulliten Mar Sci Gulf Caribb. 1958;8.

32. Lewis RJ. The Changing Face of Ciguatera. Toxicon. 2001;39: 97–106. doi:10.1016/S0041-0101(00)00161-6

33. Ledreux A, Brand H, Chinain M, Bottein MYD, Ramsdell JS. Dynamics of Ciguatoxins from Gambierdiscus polynesiensis in the Benthic Herbivore Mugil cephalus: Trophic Transfer Implications. Harmful Algae. 2014;39: 165–174. doi:10.1016/j.hal.2014.07.009

34. Loeffler CR, Robertson A, Flores Quintana HA, Silander MC, Smith TB, Olsen D. Ciguatoxin Prevalence in 4 Commercial Fish Species Along an Oceanic Exposure Gradient in the US Virgin Islands. Environ Toxicol Chem. 2018;37: 1852–1863. doi:10.1002/etc.4137

35. Randall JE. Food Habits of the Reef Fishes of the West Indies. Coral Gables: Institute of Marine Science. 1967.

